# CRISPRidentify: identification of CRISPR arrays using machine learning approach

**DOI:** 10.1101/2020.11.05.369512

**Authors:** Alexander Mitrofanov, Omer S. Alkhnbashi, Sergey A. Shmakov, Kira S. Makarova, Eugene V. Koonin, Rolf Backofen

## Abstract

CRISPR–Cas are adaptive immune systems that degrade foreign genetic elements in archaea and bacteria. In carrying out their immune functions, CRISPR– Cas systems heavily rely on RNA components. These CRISPR (cr) RNAs are repeat-spacer units that are produced by processing of pre-crRNA, the transcript of CRISPR arrays, and guide Cas protein(s) to the cognate invading nucleic acids, enabling their destruction. Several bioinformatics tools have been developed to detect CRISPR arrays based solely on DNA sequences, but all these tools employ the same strategy of looking for repetitive patterns, which might correspond to CRISPR array repeats. The identified patterns are evaluated using a fixed, built-in scoring function, and arrays exceeding a cut-off value are reported. Here, we instead introduce a data-driven approach that uses machine learning to detect and differentiate true CRISPR arrays from false ones based on several features. Our CRISPR detection tool, CRISPRidentify, performs three steps: detection, feature extraction and classification based on manually curated sets of positive and negative examples of CRISPR arrays. The identified CRISPR arrays are then reported to the user accompanied by detailed annotation. We demonstrate that our approach identifies not only previously detected CRISPR arrays, but also CRISPR array candidates not detected by other tools. Compared to other methods, our tool has a drastically reduced false positive rate. In contrast to the existing tools, our approach not only provides the user with the basic statistics on the identified CRISPR arrays but also produces a certainty score as a practical measure of the likelihood that a given genomic region is a CRISPR array.

## INTRODUCTION

CRISPR–Cas is an adaptive immune system that is present in 90%of archaeal and about 40%of bacterial genomes (1–3). These systems consist of arrays of direct repeats (CRISPR) and varying suits of CRISPR-associated (*cas*) genes that encode proteins involved in the immune functions.

The CRISPR–Cas immune response consists of three main stages:

i. Adaptation that consists of selection and excision of short target segments (protospacers) from foreign DNA followed by incorporation of these segments into the CRISPR array, to form spacers via a cut-and-paste mechanism (4–6). In most CRISPR–Cas systems, recognition of the protospacers requires the PAM (protospacer adjacent motif), a short sequence motif in the target DNA that flanks the protospacer and is crucial for avoiding self-cleavage (7–11).
ii. Biogenesis of CRISPR (cr) RNAs, including expression of the pre-crRNA (a leader followed by an array of repeat-spacer units) and subsequent processing of the transcript into mature crRNA (12,13).
iii. Interference that involves invader DNA or RNA degradation at the cognate protospacer, guided by the crRNA and catalyzed by a specific complex of Cas proteins or a single, multidomain protein (1,6,14). Typical of defense systems, CRISPR–Cas loci evolve fast and are often substantially different among otherwise closely related strains of bacteria or archaea. The CRISPR arrays are associated with *cas* genes that belong to > 50 families and form combinations specific to each type and subtype of CRISPR–Cas systems (2,3,15–18). Some CRISPR–Cas systems are associated with additional genes whose roles in defense or other functions of CRISPR–Cas remain to be established (19,20). The Cas proteins are generally encoded in single operons adjacent to the respective CRISPR array although there are many exceptions to this pattern.

CRISPR–Cas systems are currently classified into two classes, six types (I-VI) and *>* 30 subtypes. This classification is based on several criteria, primarily, the identity, co-occurrence and arrangement of the *cas* genes in CRISPR– Cas loci (2,3,18). The organization of the adaptation module is largely uniform across the diversity of the CRISPR– Cas systems. Proteins involved in adaptation are the integrase (endonuclease) Cas1, the structural subunit Cas2 and, in some systems, another endonuclease, Cas4, which facilitates the integration of PAM-compatible spacers (21). By contrast, the protein machineries involved in the crRNA biogenesis and interference are highly diverse. The two classes of CRISPR–Cas systems differ with regard to the complexity of the effector ribonucleoprotein complexes. Class 1 effectors consist of several structurally and functionally different subunits, some of which possess the pre-crRNA cleavage nuclease activity, and accommodate the crRNA to recognize the protospacer in the target DNA molecule. The target cleavage activity resides either in one of the subunits of the effector complex itself or in a separate nuclease that is recruited by the effector complex when bound to the target DNA (13,14,22,23). By contrast, Class 2 systems utilize a single, large, multidomain protein, such as Cas9, Cas12 or Cas13 that is responsible for the target recognition and cleavage and, in many case, also for the pre-crRNA processing (2,13,18). The CRISPR–Cas types and subtypes further differ in the repertoires of proteins and domains involved in pre-crRNA processing and interference. The sequence and structure of the CRISPR array itself is usually disregarded in the classification of CRISPR– Cas systems because, although CRISPR repeat organization can affect the evolution and diversity of CRISPR arrays, the relationship between repeat families and associated CRISPR–Cas types and subtypes is complicated (2,3,18,24–26). In particular, nearly identical CRISPR-repeats in the same species can be associated with different subtypes (27,28). Nevertheless, CRISPR arrays or, more precisely, the constituent repeats can also be classified into families, capturing the diversity of sequence motifs and RNA secondary structures (24–26,29–32). The existing approaches for CRISPR array identification, such as CRT (33), CRISPRCasFinder (34), PILER-CR (35), minced (https://github.com/ctSkennerton/minced) and CRISPRDetect (36), rely mainly on the repetitive structure of arrays. CRISPRCasFinder, for example, uses Vmatch (http://www.vmatch.de/) (37) to find pairs of maximal repeats, which are then joined into a consensus repeat sequence. The array defined by this consensus repeat sequence is then scored with a built-in scoring function that takes into account the lengths of the repeats and spacers, similarity between the repeats and other features. The other approaches, while differing in details, rely on a similar overall detection approach.

Here, we present a machine-learning (ML) approach for the processing and evaluation of repetitive elements in order to detect CRISPR arrays. For the repeat scoring, we do not rely on a manually curated evaluation function but rather use a ML method to learn this evaluation function from data. For this purpose, we constructed an extensive set of training and test data including both bona fide CRISPR arrays and negative controls. This allows us to substantially increase the sensitivity and to drastically improve the specificity of CRISPR array detection, while providing the user with a reliable quality measurement. The problem of false positive predictions was not thoroughly investigated in the existing CRISPR detection tools. We show that our ML-based approach has a much lower false positive rate than the other CRISPR detection methods.

## MATERIALS AND METHODS

The task of CRISPR array detection consists of the search for repetitive elements in a genome, which might form the repeats of a CRISPR array, followed by the evaluation of the identified putative array. The evaluation criteria can include the length of the repeat consensus sequence, the similarity between the repeats within the array, and features of spacers and nearby *cas* genes. The existing approaches use manually curated evaluation functions that combine these features into a score. CRISPRDetect, for example, calculates the score for an array by adding up positive and negative score contributions, such as +3 if the repeat, which is required to be longer than 23 bases, contains a known motif at the end, and a metric for identity among the spacers (−3 to +1), or −1.5 for dissimilarity between repeats. Arrays with scores above 4.0 are then classified as positive. Although such manually curated scoring functions can work well for the detection of positive examples, they have several disadvantages. First updating the scoring system to a growing database is a complicated task. Second, there is no built-in mechanism to balance sensitivity and specificity. Our analysis described here shows that, whereas the sensitivity of the existing tools is usually high, these tools do not adequately control for the false positive rate and thus have a relatively low specificity.

In order to control for both sensitivity and specificity, we set up the CRISPRidentify pipeline, which replaces the manually curated scoring function with an ML approach. The CRISPRidentify pipeline consists of two major steps, namely, a new approach to detect and generate array candidates, aiming at increased sensitivity, and a versatile, machine-learning based evaluation of the detected candidates to increase the specificity. With regard to the first task, any approach to detect CRISPR arrays faces two main subproblems: (i) accurate identification of the start and end positions for each repeat candidate in the array and (ii) accurate identification of the boundaries of the whole array. Mismatches in the repeat sequences or similarity of some spacers around the start and end positions can lead to incorrect identification of the repeat sequence itself. Incorrect identification of the array boundaries could be caused, for example, by degradation of the repeat sequence at the end of the array which makes it difficult to determine the array end point.

The second major task is the evaluation of the candidates. CRISPRidentify learns a classifier for CRISPR arrays from known data to control for the false positive rate, in contrast to the manually curated scoring functions used by the existing tools. In this setting, it is natural to rely on: (i) features that are extracted from the array for estimating the similarity between bona fide CRISPR arrays and array candidates, (ii) a benchmark set of positive and negative array examples and (iii) machine learning to learn the evaluation function from this benchmark. We define new training and test sets consisting of carefully selected positive and negative examples. Using these training sets, we train a classifier for evaluating CRISPR candidates based on 13 array-derived features (see Table S2 in Supplementary), such as the similarity between the repeats, AT-content or stability of the repeat hairpin. The principal advantage of this approach is that we not only increase the sensitivity of CRISPR array detection, but also drastically reduce the false positive rate, i.e. increase the specificity. The problem of false positive predictions has been not properly addressed by the previous approaches. Finally, the pipeline annotates the array with additional information such as its orientation, leader sequence and *cas* genes.

### Benchmark datasets

Overall, we prepared six benchmark datasets to test different aspects related to the detection of CRISPR arrays. In particular, we compiled the archaeal and bacterial arrays (29,30,38). The first dataset contained 400 archaeal arrays among which 10 were experimentally validated and the rest were manually verified (Table 1). The second dataset containing 600 bacterial arrays was constructed from published datasets (29,30,38), with the constraint that the corresponding repeat sequences should be at least 85%identical to published experimental CRISPR-repeats. For simplicity, we will refer to these datasets as archaeal and bacterial datasets, respectively. As an independent, high-quality, non-overlapping test set, we additionally selected 550 CRISPR arrays from archaeal and bacterial genomes using the following criteria: 1) the minimal number of repeats in each CRISPR array is three, and 2) the maximum distance between the CRISPR array and the *cas* gene operon is 500 nucleotides. We will call this the Cas-associated dataset (See distribution of Cas types in Supplementary materials Table S7).

**Table 1.**
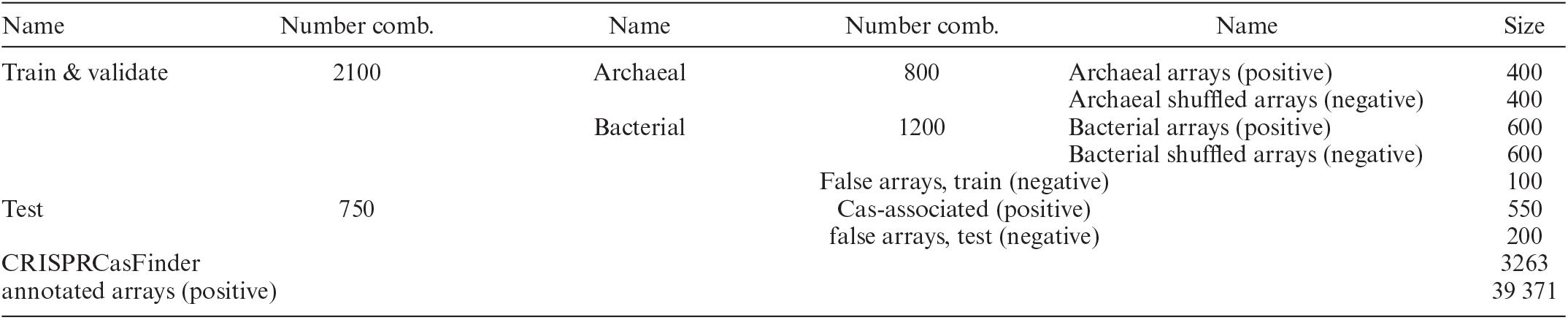
Generated data sets and their combinations used in the machine learning approach

The first three datasets were partially used by other tools, but the fourth one was designed specifically for this work and is required to control the false positive rate. It contains 300 (100 training and 200 test) false CRISPR arrays and was constructed based on the tandem repeat dataset and cases with identical ‘spacers’ (see Tables S2-S4 in Supplementary for more details). For simplicity, we will now refer to this set as false arrays. To obtain more negative examples, we also created additional distorted CRISPR arrays from the ‘archaeal dataset’ and the ‘bacterial dataset’ by independently shuffling the nucleotides in each repeat and each spacer sequences. We will refer to these as archaeal shuffled arrays and bacterial shuffled arrays, respectively. All generated datasets where split into training and test sets.

In addition to the datasets we generated, we used two datasets from literature. The first, most comprehensive one consists of all arrays annotated from the most recent CRISPR classifications (2,3,18,19), and will be called annotated arrays dataset. It consists of 39 371 arrays. The second dataset was downloaded from CRISPRCasFinder (34). It contains 3263 arrays and will be denoted as CRISPRCas-Finder dataset.

### Construction of array candidates

We use Vmatch version 2.3.0 to scan a genomic region for the occurrences of repeat candidates. Vmatch returns a list of putative repeat pairs. By default, in CRISPRidentify, we set the substring length for Vmatch to be in the range of 21 to 55 nucleotides, and the spacing between the matching pairs has to be in the range between 18 and 78 nucleotides. Both intervals can be changed by the user. To avoid duplicates, we filter out all the repeat candidates except for the unique ones.

The next step is to cluster the repeat matches found by Vmatch, and to extract the bona fide repeat sequence. As a proxy, the repeat sequence is the consensus sequence of all repeat matches belonging to a repeat cluster. However, using a consensus sequence to determine possible repeat candidates poses two problems. First, the length of the consensus sequence is uncertain because a threshold has to be defined for conserved columns, and second, we lose control over the number of mismatches to existing repeat candidate sequences in the genome. Thus, instead of using a single consensus sequence, as the first stage, we generate a set F over each Vmatch cluster containing all possible repeat candidates and their extensions. The set F is a subset of all subsequences of a maximal consensus sequence partially ordered by the subsequence relation (see Figure 1, box ‘Extension’, for an example).

**Figure 1.**
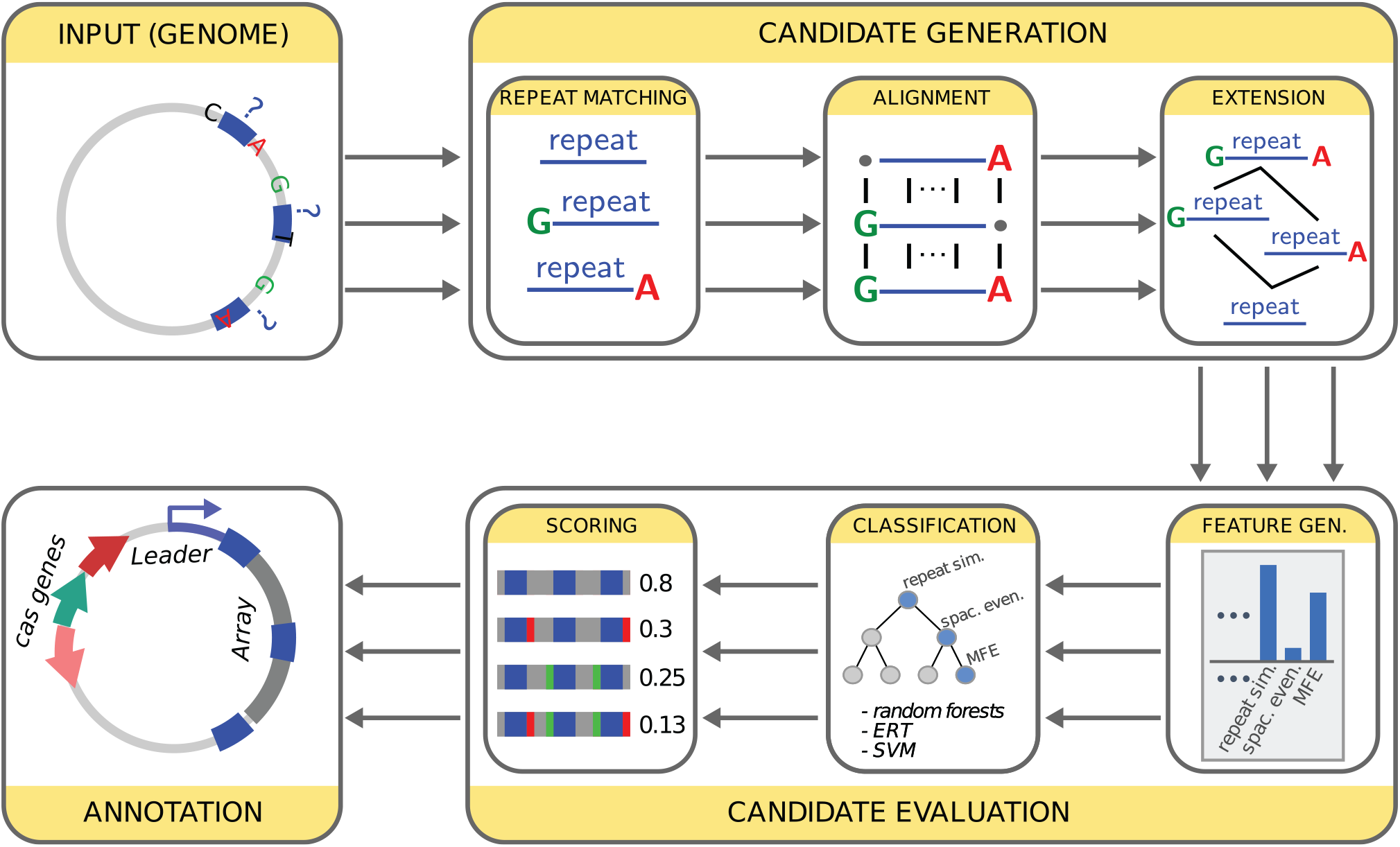
General architecture of the CRISPRidentify approach. The pipeline consists of two major steps. In the first step (Candidate Generation), candidate repeats are identified using Vmatch. Vmatch detects all potential pairwise repeats. This implies that leading and trailing nucleotides have an uncertain assignment as these repeated nucleotides could be part of the repeat, or part of a repetitive part of the spacer. For this reason, we first determine the minimal and maximal forms of the repeat by aligning the repeat candidates, and then consider all possible repeats between the minimal and maximal repeat variants. Mapping these repeat candidates back to the genome generates different array candidates, which are then evaluated using machine learning. For that purpose, in the second step (Candidate Evaluation) we extract features from the array such as repeat similarity, the evenness of spacer lengths, AT-content, minimum free energy of the repeat hairpin and others. We then use different machine learning (ML) approaches, learned on a large data sets of positive and negative examples, to classify them as CRISPR arrays. We use the scoring of the ML approaches to prioritize the different array candidates. Finally, we use different tools to annotate the array. Thus, we determine the strand orientation using CRISPRstrand and annotate the leader sequence using CRISPRleader. In addition, HMM and Prodigal to annotate *cas* genes.

To generate F, we use the alignment for each cluster to construct the maximum element of F as a string that is composed, column-wise, of the nucleotides most frequently found in the alignment. The minimum element of the set F is defined as the most consistent substring, and is a subsequence of the maximal element by definition. After the maximum and minimum elements are calculated, we generate all sequences which fulfil the requirements that they 1) are subsequences of the maximum element and 2) contain the minimal element as a subsequence. These sequences then complete the set F. Mathematically, F is a filter (the dual concept of an ideal) in the partial order defined by the subsequence relation. As the last step, the set of repeat candidates is formed via unification of sets F and initial Vmatch results and filtering of duplicates (see Supplementary Materials, Section 1 ‘E*nhancement of Vmatch results’*). Lastly, we extend the set of repeat candidates with modifications of each repeat candidate in which up to 3 nucleotides are omitted from both ends, resulting in 15 variations of each repeat. After that, duplicates are filtered out for the last time and the remaining candidates are added to the pre-existing clusters. These clustered repeat candidates are then used to form CRISPR candidates.

After the set of potential repeat candidates is extracted, we need to form the bona fide CRISPR array candidates. Unlike the other existing algorithms, where a CRISPR array is formed from a repeat consensus by simply splitting the considered interval into repeats and spacers, our approach utilizes approximate string search to find the locations of the repeat sequences using the python regex library available on PyPI regex. This approach also has an important advantage over simple partitioning of the DNA interval into repeats and spacers because this technique allows incorporating insertions and deletions as editing operations, finding the locations that minimize the number of editing operations. This feature allows our approach to effectively identify potential CRISPR candidates including cases with abnormalities, such as truncated repeat (39–41) sequences or complete spacer deletion (see section seven spacer deletion and section eight degenerated repeat in the Supplementary materials).

### Feature vector associated with an array candidate

Having obtained suitable CRISPR candidates, we generate a feature vector for each candidate to use at the evaluation stage of our pipeline. Users may specify the set of desired features by choosing one or several of the three classifier models we developed. Each of these models is defined by feature selection and encompasses a different combination of the 13 features we devised for classifying CRISPR arrays. When designing these features, we had three conceptual categories in mind, depending on whether we expect their values to be positively or negatively correlated with the likelihood of a true CRISPR array, or whether they provide supporting information. These are, however, only conceptual categories that are not further used during the machine learning approach. Among the factors, which we expect to augment the evidence for a true CRISPR array with their growing value, are number of repeats, length of the repeat sequence and repeat similarity because the more repeats are found, and the longer and more similar they are, the stronger the evidence that the result did not occur by chance.

On top of these features, we compute the RNA minimum free energy (MFE) of the consensus repeat sequence. The negative MFE is called the MFE score, and lower MFE scores are positively correlated with the likelihood of a true CRISPR array (22,27,28,42–44) in some systems. The reason is that, in many cases, for instance, in the vast majority of type I cases, the crRNA tends to form stable stem-loop structures, due to the palindromic organization of the repeat sequences. Therefore, a low MFE score can be used to identify stem-loop structures in these systems, thus buttressing the evidence for true crRNA and, by extension, CRISPR array. The MFE score was computed using RNAfold (45). Another feature positively correlated with the likelihood of a true CRISPR array is high similarity between the CRISPR candidate repeat sequence and a verified repeat sequence in the database. The similarity between a given sequence and the sequences in the database is computed using BLAST (46), which is provided as executable with our pipeline. In our approach, two BLAST similarity scores for each CRISPR candidate were computed. The first score reflects the highest similarity between a candidate and all experimentally validated repeat sequences. The second score measures the similarity to a modified version of the same dataset to which one or two mismatches were added, thus extending the range of similarity considered by the first BLAST score. These two scores were treated as two separate features.

In contrast to the first group of features, the values of the following features are negatively correlated with the likelihood that a candidate is a true CRISPR array. The first such feature is the number of mismatches because repeats are usually highly conserved. Furthermore, spacers of the CRISPR array candidate were assessed in terms of length and similarity. Because in bona fide CRISPR arrays, the spacer length is typically uniform, a high variance in spacer length negatively correlates with the likelihood of a CRISPR array. With respect to similarity, spacers differ from repeats in that they are generally not significantly similar to each other given their origin from different virus or plasmid genomes. Thus, high level of similarity between spacers (see Supplementary Figure S2 in Supplementary) is negatively correlated with the likelihood of a candidate being a CRISPR array. Another feature that is included in our pipeline is whether a candidate region contains a protein-coding sequence. Because CRISPR arrays do not code for proteins, the presence of coding sequences is negatively correlated with the likelihood of an interval being a CRISPR array. Our pipeline identifies Open Reading Frames (ORFs) using Prodigal (47), again provided as executable with our pipeline. If ORFs were found, the result with the highest confidence value was taken as the feature value. To account for protein coding-regions even more thoroughly, our tool also looks for tandem repeat proteins. To this end, the Prodigal output is fed into a Hidden Markov Model (48), which searches for similarity with the PFAM database of tandem repeat proteins. The resulting tandem protein similarity score serves as a feature negatively correlated with the likelihood of the interval being a CRISPR array.

In the third category, our approach computes two features whose growing value is not directly correlated with the likelihood of the interval being a CRISPR array but could potentially guide the likelihood in different ways. The first such feature is the AT-content of the identified repeat sequences within a given CRISPR array candidate. Because bacteria and archaea differ in AT-content, this feature can help to define implicit subclasses that allow the classifier to model different properties for bacterial and archaeal arrays. The other additional feature is average spacer length (see Supplementary Figure S1 in the Supplementary Materials). This feature is important because, in contrast to repeats, spacers in the spacer array can have different lengths. Overly short or long average spacer length, however, might signal a false array, which is why such outliers should be considered in the scoring of the candidates. After all the features determined by the classification model were calculated for each CRISPR array candidate, the resulting up to 13-dimensional feature vector was subject to machine learning classification as described below.

### Feature selection

After the first successful results using all thirteen features described above, we went on to evaluate which of the features are actually necessary. This can be determined by feature subset selection (FSS), which generates a subset of features best suited for classification. For this purpose, we applied a wrapper approach (49), which works as follows. We generated all 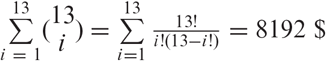 subsets of the full sets of 13 features. We then trained for each of the 8192 subsets a model using only the features from the respective subset, and assessed the prediction quality. The results of these experiments showed that a subset of the 13-dimensional feature vector is sufficient to reach the 99.8%accuracy that is achieved with the complete feature set. The same accuracy levels could be achieved with three of these models. These models included 8, 9 and 10 features, respectively, with each feature appearing in at least one of the models. Although the feature combinations overlapped, they were not subsets of one another (see Table S1 in Supplementary for the models and their feature subsets).

### Classification of CRISPR arrays using machine learning

The last step of our CRISPR identification pipeline consists of the evaluation of the obtained CRISPR candidates. For this purpose, we integrated the implementation of the Extra trees classifier from the python Scikit-learn package (50) into our pipeline. We provide the user with an opportunity to pick either of the three pretrained classifiers that we describe in the Feature Selection and Importance Analysis subsection or use all of them simultaneously. It is important to notice that we do not use the class prediction directly but take the label pseudo-probability and treat it as a certainty score. In case of using many classifiers, the score is computed as the average of scores. Based on the distribution of the output CRISPR array scores, we introduced two decisive score values around which we structured the labels: 0.75 and 0.4. CRISPR arrays with scores greater than or equal to 0.75 were labeled to ‘candidate’, meaning that one of them likely corresponds to a true CRISPR array. CRISPR arrays with scores between 0.75 and 0.4 were assigned to the group ‘potential candidate’, meaning that there is a smaller probability for one of them to be a true CRISPR array. The CRISPR candidates with scores lower than 0.4 were assigned to the group ‘low score candidates’, indicating that they are probably not valid CRISPR arrays. After performing the grouping, the algorithm picks the ‘bona fide candidate’ as the candidate with the highest score among the candidates in the ‘possible candidate’ group. Then all the candidates are reported in files corresponding to their groups. Hence, the user may not only inspect the bona fide candidates but also all the alternatives with their respective statistics.

### Repeat and spacer databases

Our tool can also be used to build a user-specific CRISPR array databases. In order to illustrate the potential of this feature, we used data from archaea and bacteria genomes. The archaeal dataset consisted of 251 complete and 736 incomplete genomes, while the bacterial dataset included 1693 complete genomes and 25335 incomplete ones. Having identified CRISPR arrays in each dataset with our tool, we extracted the complete set of spacers as well as the consensus repeat sequence for each CRISPR array. Using these results, we built two databases, one for the identified repeats and the other one for spacers. The output of the repeat database lists, for each consensus repeat sequence, all genomic locations, the organisms, as well as the number of locations and organisms where an array with this consensus repeat can be found (see Table S6 and Figure S6 in Supplementary).

## RESULTS AND DISCUSSION

### Existing CRISPR datasets contain many suspicious arrays

The main drawback of the current tools for CRISPR array identification is the lack of scoring of the obtained results, i.e. absence of a thorough evaluation of the likelihood of each reported array to be a true CRISPR array. We suspected that lacking a stringent quality evaluation might enrich existing datasets with false-positives. A high prevalence of questionable results produced by the commonly utilized CRISPR identification tools could pose a problem not only to experimental researchers, which might invest effort in the investigation of false predictions, but also to tool developer using these datasets as standards (51).

One type of spurious arrays that we found to be common in public databases are arrays that contain damaged repeats or spacers with unusually high similarity to one another. In these cases, a possible explanation is a duplication or repetitive structure stemming from a source other than CRISPR. Consider the example in Figure 2, which is an array that was predicted as positive by the CRT tool. As shown in Figure 2A, the array contains a damaged repeat as well as two nearly identical spacers that differ by only one nucleotide. Further examination showed that the array lies within a predicted protein-coding sequence on the opposite strand, suggesting that this is not a functional array. Moreover, the CRISPR locus is in a region that is predicted to be intrinsically disordered in the protein encoded on the opposite strand, and the repeat (and part of the spacers) are predicted by the Anchor tool (52) to be linear interacting peptides (Figure 2B). Thus, in this case, the repetitive structure at the nucleotide level, most likely, originates from a repetitive structure at the protein level, and accordingly, is not a functional CRISPR array. Other suspicious cases showing, for example, low repeat similarity or a highly non-uniform distribution of spacer lengths can be found in Section 9 ‘*Spurious Arrays in Existing Tools’* and Supplementary Figures S3–S5 of the Supplementary material.

**Figure 2.**
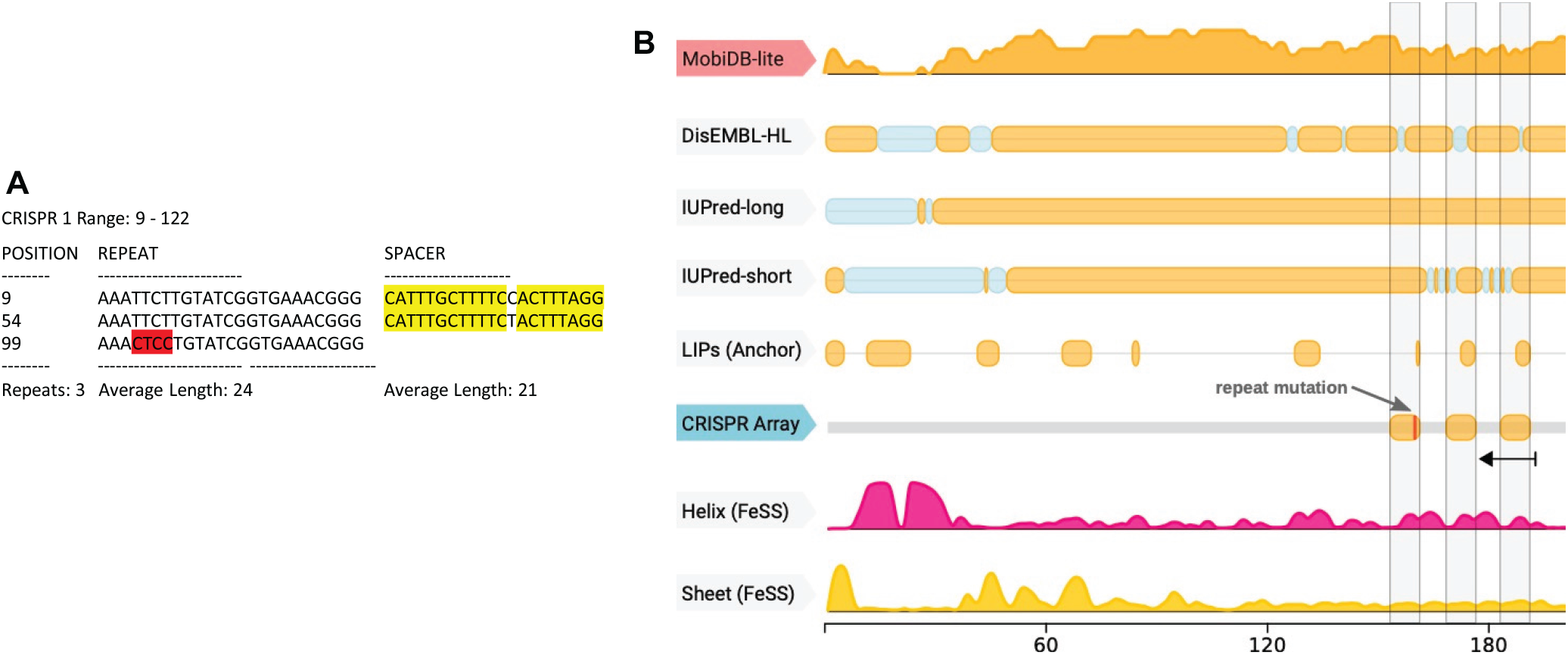
Example of a spurious detected array. (**A**) The repeat spacer structure is special in two aspects. First, the two spacers are identical (marked in yellow) with the exception of one position. Second, the third repeat has several mismatches (marked in red). (**B**) The locus is overlapping a predicted protein on the opposite strand, which has an intrinsically disordered region predicted by several tools to overlap the array position. Furthermore, the three repeats overall exactly with three linear interacting peptides (LIPs) predicted by Anchor. In addition, the first LIP is terminated at the position where the last repeat (in opposite direction) has a mutated region. This together provides a repetitive protein sequence or a duplication event as an alternative explanation for the repetitive structure predicted as CRISPR array.

To assess the scale of the problem, we investigated the 3263 array candidates in the CRISPRidentify benchmark set and systematically searched for suspicious arrays that are predicted as positive by CRT, CRISPRDetect and CRISPRCasFinder. We considered an array to be suspicious if it had at least one of the following properties: (i) a high similarity between spacers (sequence identity > 0.6), (ii) mutations in repeats (identity between repeats *<* 0.8), (iii) similarity between repeat and spacer (identity between repeat and spacer > 0.5) or (iv) an uneven spacer length distribution. The spacer length was considered to be distributed unevenly if at least one spacer was 70%shorter than the average. To exclude any bias in the tool parameter setting, we considered only arrays that could be handled by all tools. Thus, we filtered out the cases with only two repeats, or arrays with repeats shorter than 23 or longer than 55 nucleotides as this setting is covered by all tools. Note that each tool predicted slightly different locations overlapping these arrays. Altogether, we obtained 1358 arrays that passed this filter and were predicted as positive by CRT, 1395 by CRISPRCasFinder and 346 by CRISPRDetect. Among these predictions, 602 arrays predicted by CRT, 628 arrays predicted by CRISPRCasFinder and 110 arrays predicted by CRISPRDetect were deemed suspicious under the above criteria (see Table 3). This corresponds to potential false positive rates of 44.3%(CRT), 45.0%(CRISPRCas-Finder) and 31.8%(CRISPRDetect) (see Table S5 in Supplementary). Although these numbers are upper bounds because it cannot be ruled out that a suspicious array is actually functional, they clearly show that false positive predictions are a substantial problem for all available methods for CRISPR detection.

### The CRISPRidentify Pipeline using a machine-learning based scoring system

To overcome the problem of false positives, we set up the CRISPRidentify pipeline (see Figure 1). It consists of two steps, namely, a sensitive approach to detect array candidates, and rigorous evaluation of the detected candidates using machine learning.

In the first step, we used the Vmatch tool to detect pairwise matches of a repeat, from which we reconstruct array candidates by combining compatible pairwise repeat matches. Vmatch builds an efficient computational index structure named suffix arrays that allows for a fast search for matching substrings in the DNA sequence. Pairwise matches that fulfill predetermined length and spacing requirements based on reliable observations are then processed further (see Methods for details). The principal difficulty at this step is that assignment of the leading and trailing nucleotides of the spacers can be ambiguous. Thus, mismatches in the repeat sequences or similarity of some spacers around the start and end positions can lead to incorrect identification of the repeat sequence itself. Consider the following repeat composition:

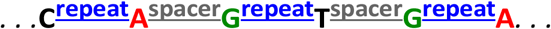

It remains unclear whether the G- and A-nucleotides coloured in green and red, respectively, belong to the repeats, or to the spacers that happen to contain the same first (or, respectively, last) nucleotides in two cases. Repeat finding tools, such as Vmatch, find pairwise matches, which are likely to be copies of the same repeat. Thus, in our example, Vmatch would return the following pairwise matches that could all be instances of the repeat: 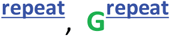 and 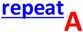 This situation results in three distinct potential array variants. To identify these cases, we first cluster the pairwise matches representing repeat candidate sequences based on the distance between them. However, the correct repeat candidate might not be included in the Vmatch results due to potential similarities in the beginnings or ends of spacers, or mismatches between repeat sequences. In the simplified example above, the candidate 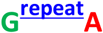 would not have been found as an exact pair match because there is no second example of this form in the genome. To overcome this problem, we extend each cluster in an enhancement procedure by additional repeat candidates. First, we generate a set F over each Vmatch cluster containing all possible repeat candidates and their extensions. To generate F, we use the alignment for each cluster to construct the maximum element of F as a string that is composed, column-wise, of the nucleotides most frequently found in the alignment. The minimum element of the set F is defined as the most consistent substring, and is a subsequence of the maximal element by definition. The set F is a subset of all subsequences of a maximal consensus sequence partially ordered by the subsequence relation (see Figure 1, box candidate extension). Mathematically, F is a filter in the partial order defined by the subsequence relation. It contains all subsequences that contain the minimal repeat element and are itself subsequences of the maximal element (see Methods and Figure 1, box titled ‘Extension’, for details).

As a result, the first step of our pipeline produced multiple CRISPR array candidates for the same genomic region. Thus, an essential second step is to evaluate all these candidates under stringent criteria. On the one hand, this evaluation guarantees that the best candidate is selected for each genomic region. On the other hand, the task of the evaluation is to exclude repetitive patterns that might be detected as CRISPR arrays by other tools but show otherwise little similarity with bona fide CRISPR arrays, resulting in a better control over the false positive rate. For this purpose, it appears natural to rely on 1) a benchmark set of positive and negative array examples, 2) features that are extracted from the array to estimate the similarity between bona fide CRISPR arrays and array candidates, and 3) machine learning to learn the evaluation function from this benchmark.

The positive examples for the benchmark set are easily generated by selecting an appropriate subset of annotated CRISPR arrays (see Materials and Methods). The reconstruction of negative examples, however, is more challenging as there is no curated benchmark set for this purpose. Here, we constructed a negative data set based on the tandem repeat dataset and cases with identical ‘spacers’ (see Materials and Methods for more details).

For the purpose of ML-based evaluation, our pipeline associates to each array candidate a feature vector, which is a set of numerical values (features) that describe different properties of the array. In our pipeline, the feature vector consists of a maximum of 13 features, including repeat-associated features, namely, number of repeats, length of the repeat sequence, repeat similarity, number of mismatches, AT-content, and the RNA minimum free energy (MFE) of the repeat, which is indicative of a stem-like structure present in the vast majority of type I arrays. We added also added two BLAST scores that determine the similarity of the identified repeats to known repeat sequences. Spacer-related features are similarity between spacers, average spacer length and spacer uniformity, i.e. a measure of the variability of the spacer length. Additionally, we included two features that are associated with the whole array, namely, an ORF score, which measures whether the array overlaps with a protein-coding region, and a tandem protein similarity score, which is used to detect tandem repeat proteins (see Materials and Methods and Table S2 in Supplementary for details). Once the feature vector was associated with a repeat region, a trained classifier was used to decide whether an array candidate is likely a CRISPR array, and the score for this likelihood was calculated.

CRISPRidentify not only defines and scores CRISPR arrays, but provides the user with more detailed information on the respective CRISPR–Cas systems for further investigation. Thus, we use CRISPRstrand (30) to predict the orientation of the array, and CRISPRleader (38) to predict the start of the leader sequence. In addition, the results of the CRISPR array analysis are complemented with the information on the detected adjacent *cas* genes. We used Prodigal (50) for the protein prediction, with subsequent classification using Cas-specific Hidden-Markov-Models (HMM). An important aspect of our annotation is the analysis of degenerated repeat elements, as well as detection of insertion sequence (IS) elements, a type of transposable elements that can influence the prediction of CRISPR arrays. A degenerated repeat element can be missed during the CRISPR array identification due to a high number of mismatches exceeding the detection threshold. To overcome this problem, we apply an additional search for degenerated repeat sequences using the approximate search method from the CRISPR identification part.

The IS elements are widespread in archaeal and bacterial genomes (53) and thus can disrupt CRISPR arrays. Integration of an IS element into a CRISPR array would mislead existing CRISPR detection tools to predict two or more distinct CRISPR arrays, despite the relatively close positioning and identity of the repeat sequences. To detect IS-elements, the known IS sequences were clustered using the Markov Cluster Algorithm (MCL) (54) and an HMM model for each cluster using hmmbuild (48). When two neighboring CRISPR arrays with identical repeat sequences were detected, Prodigal was used to extract protein sequences from the gap region, and the resulting sequences were searched for the presence of an IS-element using the IS-HMM models. If an IS-element is found, our pipeline has an option to merge the separated parts of the array (see the *Degenerated repeat and IS element search* section in the Supplementary Materials for an example).

### CRISPRidentify outperforms the existing tools thanks to machine-learning based evaluation

We investigated different types of machine learning (ML) approaches, including Support Vector Machine, Naive Bayes, K-nearest neighbors, Fully Connected Neural Network, Decision Tree, Random Forest classifier and Extra-trees classifier, for the use in CRISPRidentify. The performance analysis for the CRISPRidentify pipeline consists of two parts. First, we determine which of the ML-approaches performs best by checking the classification performance of each. Then, we compare the selected, best ML-approach to the existing tools, using the built-in scoring function and thresholds of these tools as the classifier. To avoid any bias from the training in the comparison with other tools, we split the positive and negative datasets into a train&validate dataset to determine the best ML-approach and parameters for CRISPRidentify, and an independent test dataset that was only used for the comparison to other tools (see Table 1 and Methods for details on benchmark sets). With the train&validate dataset, we used a 10-fold cross-validation strategy to assess the accuracy of the different ML-approaches. As the train&validate dataset is balanced (see Table 1), reporting the accuracy is sufficient to assess the performance of each approach. As can be seen in Figure 3, ensemble methods, in particular, Extra Trees, which learn a whole ensemble of simple classifiers and output the average estimate for all of them as the overall classification result, clearly outperform the other classifiers with a median accuracy of 0.91. Thus, we employed Extra Trees for the further analyses.

**Figure 3.**
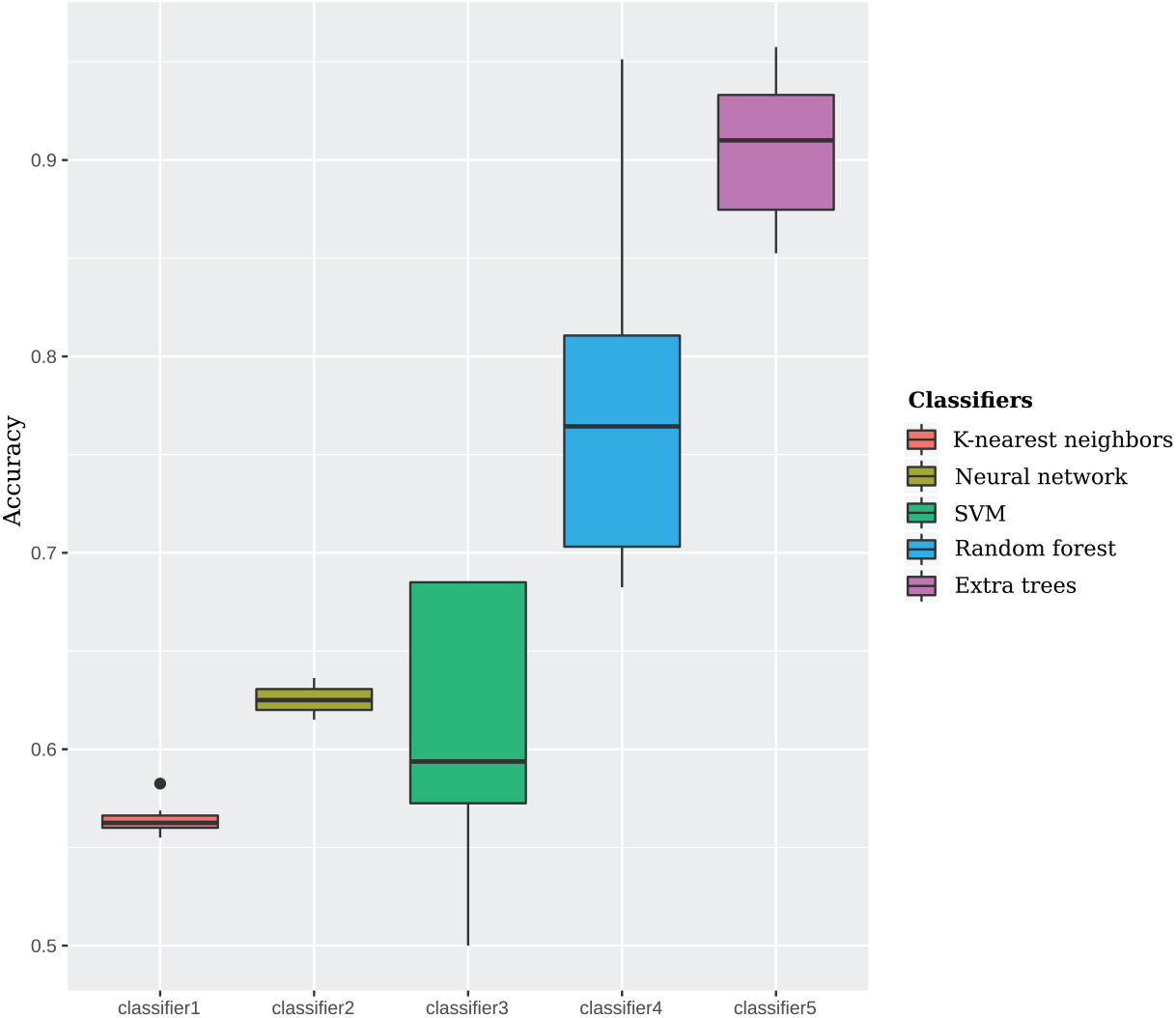
Performance of different Machine Learning classifiers. We conducted the grid search of hyperparameters for each of the classifiers (See supplementary Materials Section 13). The initial experiment shows the dominance of the ensemble methods for CRISPR array classification. See the variety of *cas* gene types in the Supplementary Materials Section 14

Once we have selected and trained Extra Trees on the train&validate set, we used the untouched **test** set (see Table 1) to compare the performance of our method using the chosen classifier with those of other tools. Because our test set is unbalanced due to the smaller amount of negative data, we report, in addition to the accuracy (ACC), also the balanced accuracy (BACC) and the Mathews correlation coefficient (MCC, see Figure 3 and Table 2), as these prediction quality measurements are better suited in this case. The high true positive rate (TPR) demonstrates the high sensitivity of all approaches, with CRT having the lowest sensitivity (TPR = 0.93) and our tool, CRISPRidentify, having the highest sensitivity (TPR = 0.99). The specificity, however, is drastically different between CRISPRidentify and the existing tools. As indicated above, means to reduce the false positive rate (or equivalently, increase the true negative rate, as FPR = 1 – TNR) was not investigated in great depth for the existing tools. Thus, CRISPRidentify has the highest possible TNR of 1, whereas CRT has a low TNR of 0.42, and CRISPRCasFinder and CRISPRDetect show intermediate specificities (TNR of 0.64 and 0.74, respectively). The relatively low specificity of the other tools is also manifested in their low balanced accuracy and Mathews correlation coefficient.

**Table 2.**
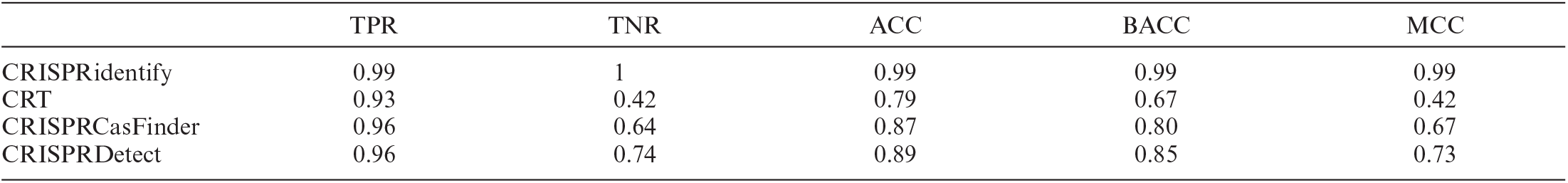
Performance of CRISPR detection tools on the test dataset, which was not used for training CRISPRidentify. As the test set is imbalanced, we report, beside the accuracy, also the balanced accuracy and the Matthews correlation coefficient

**Table 3.** Spurious Arrays in the CRISPRCasFinder datasets (27). To detect arrays that are likely false positive candidates, we first filtered arrays that are not covered by the parameter setting of the different tools (i.e. ≤ 3 repeats, or repeat length < 23 or >55). The remaining number of arrays is listed in the row *standard arrays*. Arrays that were labeled as *likely negative* (see Methods) if they had one of the following four defects: 1) damaged repeats, 2) high similarity between spacers, 3) uneven distribution of spacer lengths and 4) high similarity between repeats and spacers. The following rows list how many of these likely negative arrays had specific deficiencies (see text for details). Note that some arrays can have several problems, hence there are multiple entries in these rows. The FDR is calculated considering all likely negatives as true negatives, as we do not have other information. This FDR values are, however, likely overestimates. Nevertheless, they show the dimension of the problem

|  | CRT | CRISPRCasFinder | Detect |
| --- | --- | --- | --- |
| Arrays declared positive | 1866 | 3263 | 764 |
| Standard arrays | 1358 | 1395 | 346 |
| Likely negative arrays | 602 | 628 | 110 |
| of which are arrays |  |  |  |
| with damaged repeats | 165 | 189 | 40 |
| with similar spacers | 57 | 0 | 13 |
| with uneven spacers lengths | 473 | 508 | 83 |
| with sim. repeats to spacers | 23 | 4 | 11 |
| FDR | 44,3% | 45,0% | 31,8% |
| likely positive arrays | 756 | 767 | 236 |

We then investigated the false positives in more detail to see whether a particular type of arrays was preferentially misclassified by a group of tools. Examination of the overlap between different tools shows that the majority of the false positives are unique for a specific tool (see Figure 4). From the 200 false arrays of our test set, all have been misclassified (that is, recognized as CRISPR) by at least one of the existing tools, and none by CRISPRidentify. A substantial majority of the false arrays (159, or 79.5%) were mis-classified by exactly one tool, and only one array was mis-classified by all three tools. In addition to excluding a specific bias, these observations support our selection of false arrays as true negatives. Among the tested tools, CRT had the highest false positive rate (117 false positive predictions (FP), FPR = 58.5%), followed by CRISPRCasFinder (72 FP, FPR = 36%) and CRISPRDetect (53 FP, FPR = 26.5%) (See feature distribution of false predictions in Supplementary Figures S12–S16). The CRISPRidentify had, as stated before, a zero false positive rate.

**Figure 4.**
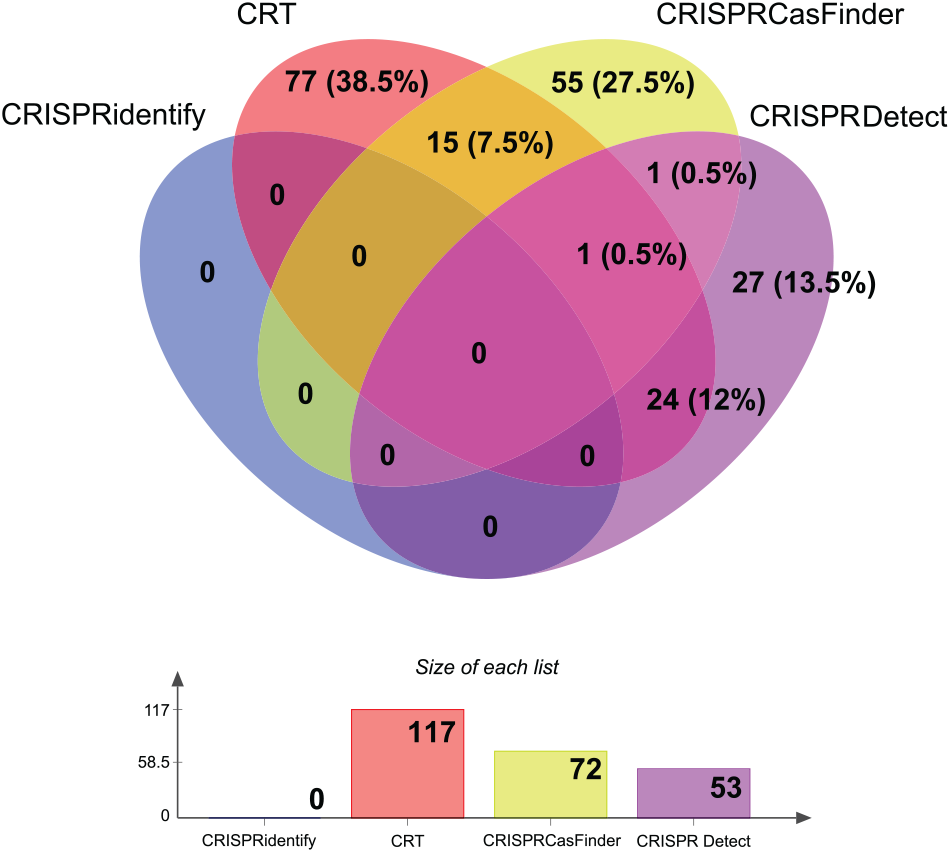
Comparison of false positive predictions on the test dataset with 200 false arrays. In summary, all false arrays were reported as a true array by at least one existing tool. In contrast, CRISPRidentify predicts all 200 as true negatives. Even further, the majority (159 out of 200) of the arrays were reported as a true array by only one approach, validating the quality of the false data set all these 159 arrays where reported as false arrays by two existing tools. For more details about the datasets see corresponding statistics in the Supplementary Table S4 Section 10. Corresponding distribution plots are presented in Supplementary Section 19.

We then investigated the true positive predictions, i.e., correctly reported CRISPR arrays, as well as false negative predictions, in more detail. We employed the full, comprehensive dataset of all annotated arrays from the most recent CRISPR classifications (2,3,18,19) (see Materials and Methods) to obtain a wider picture of arrays that remained undetected by different tools. This dataset or a larger part of it were not used for training the parameters for several reasons: (i) the dataset would be highly imbalanced, (ii) we would be unable to perform the same quality control, due to the dataset size, as for the train&validate dataset and (iii) the arrays are probably largely predicted with one of the existing tools. The last point is problematic as it would bias our tool to mimic the prediction tool that was mostly used to annotate these arrays. As shown in Figure 5, the sensitivity is high for all existing tools, but nevertheless, CRISPRidentify performed the best (99.8%), followed by CRISPRCas-Finder (98.9%), CRISPRDetect (94.5%) and CRT (93.5%). In this test, CRISPRidentify could detect 147 previously annotated arrays (2,3,18,19) that were not detected by any other approach. Overall, only 89 arrays, 7%or the total of 35310, were predicted by all tools, indicating considerable variation among the predictions. In summary, these results show that, in CRISPRidentify, we increased the already high sensitivity of the existing approaches, while sub-stantially improving the specificity as well.

**Figure 5.**
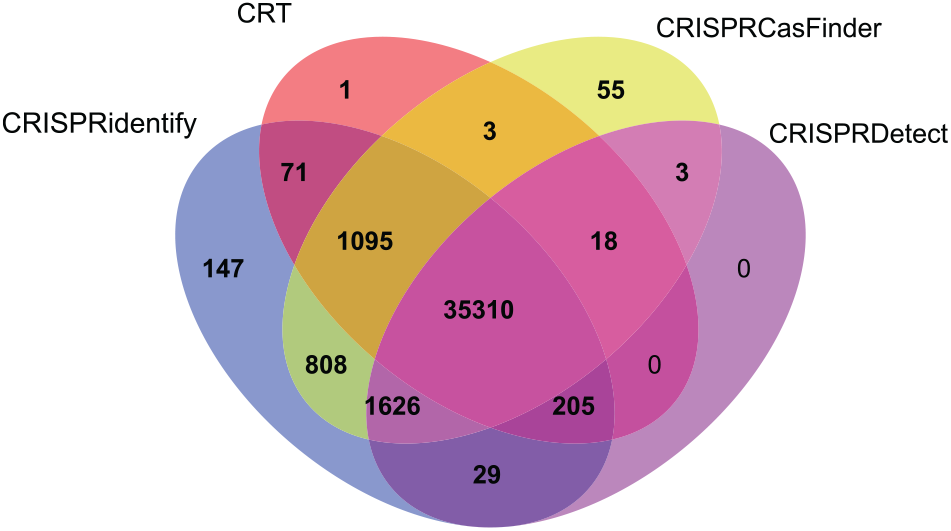
CRISPRidentify recall level in comparison with other tools on the annotated arrays dataset. This dataset was compiled out of 987 archaeal and 27 028 bacterial genomes obtained from NCBI. Only 89, 7%of the annotated arrays, are predicted by all tools. 147 arrays are only predicted by CRISPRidentify, which is nearly three times as many as the 55 arrays predicted only by CRISPRCasFinder. Concerning the largest overlap between tools outside the commonly predicted 35310 arrays, 9.0%or 808 + 1095 + 3 arrays are predicted by both CRISPRCasFinder and CRISPRidentify.

### Both expected and unexpected features are important for CRISPR identification

To assess the importance of different features, we first performed feature selection to remove non-informative features. This procedure also helps avoid overfitting to the training data. Thus, we generated three subsets of features including 8, 9 and 10 features, respectively, that are sufficient to reach the 99.8%accuracy. Although the feature combinations overlapped, they were not subsets of one another (see Methods for details, and Table S1 in Supplementary for the models and their feature subsets).

We then investigated how much each feature contributes to the prediction quality. Specifically, we used the ReBATE algorithm since it showed state of the art performance for the similar task (55). We ran the algorithm on the training set with different numbers of the nearest neighbors within the interval [1, 100], to compute the relative importance of each feature. We then compared the importance values obtained with the training set with the feature importance of the trained model. To this end, we sampled 10 000 CRISPR candidates from the comprehensive dataset and re-ran the ReBATE algorithm with the same settings. Despite minor differences, both runs showed similar results with slightly different orderings of spacer similarity, spacer uniformity and AT-content (see Figure 6).

**Figure 6.**
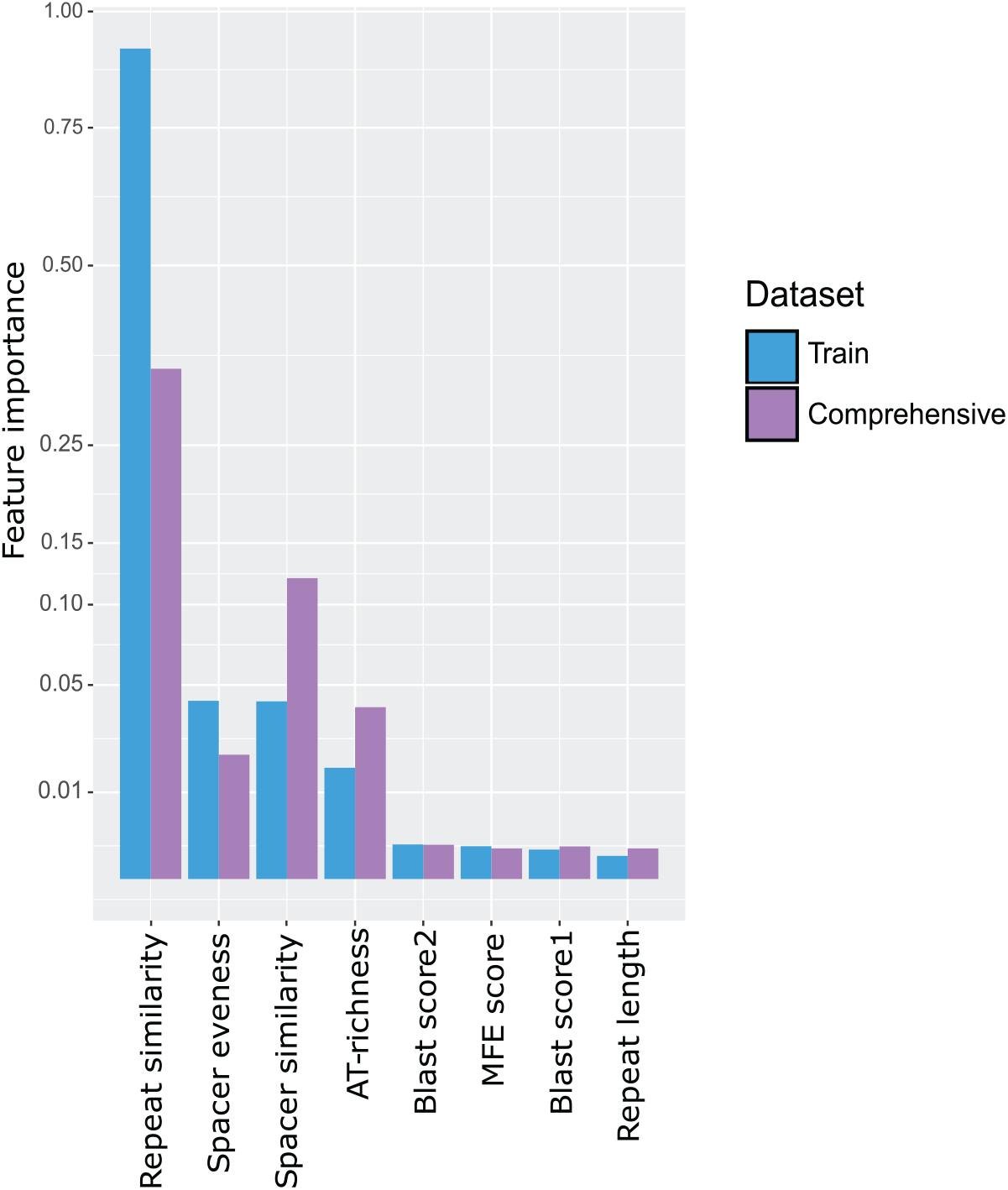
Feature importance distribution of eight most influential features which was computed with ReBATE algorithm. Feature importance distribution on the train dataset described in Table 1 and sampled cases from the comprehensive dataset classified with our approach. The sampled results contained 10 000 classified CRISPR array candidates.

Overall, as could be expected, repeat similarity is clearly the most important feature. The MFE-feature is also expected to contribute substantially because the repeat forms a stem-loop structure that is required for processing in the vast majority of type I systems. The AT-content did not come across as a direct decision-making feature, i.e., there was no correlation between AT-content and array quality. Thus, this appears to be a supporting feature defining sub-classes that can help the classifier to compensate for the differences between archaeal and bacterial arrays, probably, also in combination with the MFE feature. The blast score is important because similarity of the repeat sequence to a known repeat is a clear sign of a true array. The importance of this feature, however, is not high enough to jeopardize the generalization capacity of the trained model. Repeat length is also an important feature as the majority of known arrays in bacteria show a narrow distribution of the repeat length, with peaks around 28 and 36 nt, and in archaea, there are three peaks of repeat lengths (Figure 7). We also investigated the distribution the repeat lengths in the false arrays, finding that this distribution is somewhat broader than the distribution for the bona fide arrays, but covers a similar length range (Figure 7). This also shows that our set of false arrays shares basic properties with the true ones.

**Figure 7.**
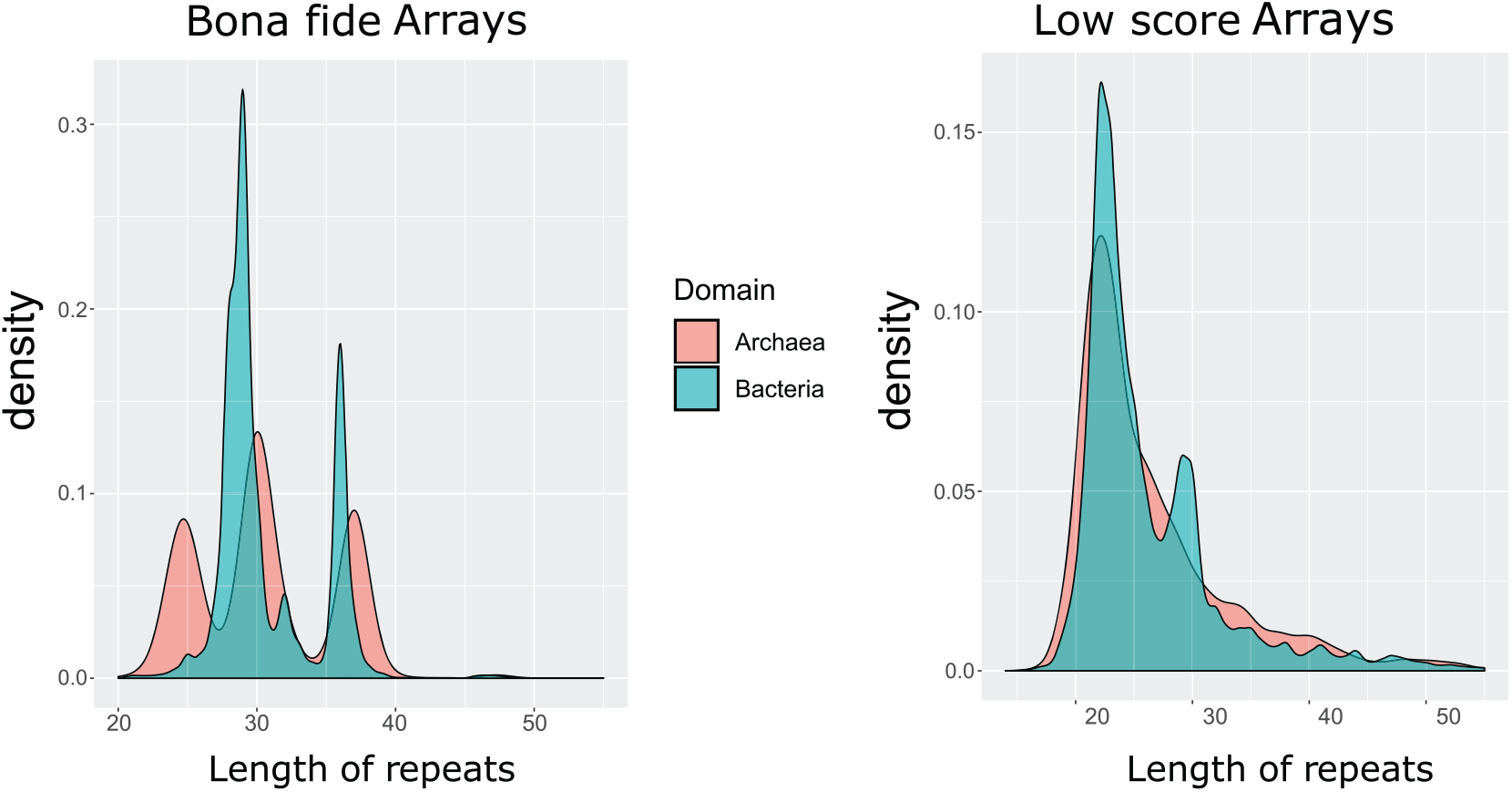
Distribution of repeat lengths in archaea and bacteria for positive and negative arrays. CRISPR arrays were obtained by CRISPRidentify on the **annotated arrays** dataset (see Table 1. *Bona fide Arrays* were defined as candidates that obtained certainty score (probability of being a CRISPR array) higher or equal to 0.75, *Low score Arrays* obtained certainty scores lower than 0.4). See separate distributions for archaea and bacteria in Supplementary Section 18 Distributions of arrays with different scores.

Whereas the importance of all repeat-related features could be readily expected, we found that, among all features, the second and third most important ones (second and fourth in the comprehensive data set, see Figure 6) are related to the spacers. Spacer similarity was one of the criteria used for constructing the negative set and is thus negatively correlated with the likelihood of the candidate being a true CRISPR array. The reason is simply that the presence of many (nearly) identical spacers in an array is strong evidence of a non-CRISPR repetitive element. However, we observed extensive overlap between spacer similarity in the positive and negative training set, showing that spacer similarity alone is not sufficient to discard an array (see Supplementary Section 3). This conclusion was supported by database analysis in which we found a few cases of spacer duplication in an array. Further, we found that the spacer lengths have a sharp peak around 30–35 for bacteria (see Supplementary Section 3), which is why spacer uniformity is an important feature as well.

In addition, we used the comprehensive dataset (see Supplementary material Section 12) for a quantitative comparison of bona fide CRISPR arrays and genomic regions that exhibit similarities to CRISPR arrays but are likely to be true negatives, that is, low scoring candidates detected by our pipeline. We found that such low scoring candidates tend to have much shorter repeat sequences, with the majority having a repeat length of about 20 nucleotides. In contrast, the vast majority of bona fide arrays had longer repeats consisting of more than 30 nucleotides. Another striking difference pertained to the number of repeats per array. Whereas low score candidates consistently contained 3–5 repeats per array, bone fide arrays were distributed more uniformly, often showing cases with large (>30) number of repeats. Further, the spacer length distribution of bona fide arrays was heavily concentrated within the interval between 27 and 50 nucleotides. Low scoring candidates, in contrast, showed a rather uniform distribution of spacer lengths between 0 and 60 nucleotides (see Figures S7–S11 in Supplementary materials)

### Repeat and spacer databases

In addition to CRISPR array identification, our tool also includes a feature that allows users to build databases that collate relevant information on the identified arrays. The advantage of this database is that it classifies CRISPR arrays into different categories (bona fide, alternative, possible, low score) and provides detailed information on each array, such as the repeat sequence, the spacer elements, the array score, its orientation, and other characteristics (see Materials and Methods for details).

### CRISPRidentify can handle common challenges faced by existing tools

Multiple studies show that, in many functionally characterized CRISPR arrays, there is notable accumulation of point mutations in the repeats at the 3’-end (41,56,57). To assess the frequency of such mutations, we investigate the bona fide arrays obtained by CRISPRidentify in a comprehensive genome dataset (see Supplementary Section 12). We filtered out cases with an overall repeat similarity below 0.8 and repeat number < 3, in order to focus on readily identifiable substitutions in the first or last repeat. The array orientation was determined using CRISPRstrand. In an effort to attain an even higher specificity, we introduced two groups of mutated terminal repeats: (i) damaged, where the number of substitutions in the first or last repeat of the array did not exceed 5 and (ii) severely damaged (more than five mismatches with the consensus repeat sequence), where we suspect degenerated repeat sequences in the array (39–41) (here, using CRISPRidentify, we found that substitutions in 3’-end repeats were frequent, appearing in 10 425 of the 39 378 analyzed arrays (27%). We also noticed that severe damage occurred in nearly a half of the mutated repeats, 5125 (13%) (See Supplementary materials Table S8). We examined how the existing tools could handle the identification of the repeats with mismatches by checking for the predicted start and end positions for each array. Of the arrays with mismatches detected by CRISPRidentify, CRT, CRISPRCasFinder and CRISPRDetect reported 7974 (76%), 8523 (82%) and 387 (4%) cases in the damaged category, respectively, and 3518 (68%), 3714 (72%) and 8 (0.1%) cases in the severely damaged group, respectively. While CRT and CRISPRCasFinder yielded similar results, CRISPRDetect reported many fewer cases. This difference is due to the conservative approach adopted by CRISPRDetect. Despite its ability of tolerate mismatches, CRISPRDetect often does not include the damaged repeat in the candidate set. In contrast, CRT and CRISPRCas-Finder can both add heavily damaged repeat candidates in the predicted arrays (see Supplementary Section 16).

Another potential challenge for CRISPR array identification could be the presence of identical spacers in the array. In order to investigate this case, we examined bona fide CRISPR arrays obtained by CRISPRidentify from a comprehensive genome dataset (see Supplementary Section 12) as well as CRISPR arrays from the training and test sets for the classification. We took into account two possible configurations: first, a sequence of identical spacers forming a ‘cluster’ of consecutive elements, and second, sparse identical spacers separated by other, distinct spacers. For the first case, our tool identified 2391 arrays, and for the second case, 1444 arrays of the total 39 378 arrays analyzed (see Table S9 in Supplementary materials section 18, also see Tables S10 and S11 for the analogous distribution on the train and test sets). The majority of the identified arrays with identical spacers contained less than three spacer duplicates. This result implies that, although our approach penalizes high similarity between spacers, it nevertheless can detect arrays with a limited number of repeated spacers, while discarding those in which many spacers are identical. In contrast, the existing tools do not evaluate the detected CRISPR arrays for spacer similarity and therefore can report spurious candidates with large clusters of identical spacers. Indeed, we examined the results of CRT, CRISPRCasFinder and CRISPRDetect, and found that all these tools reported some spurious CRISPR array candidates with large clusters of identical spacers (*>*5) (see Supplementary Section 18). Finally, our approach is able to identify CRISPR array candidates with complete spacer deletion, in contrast, to CRT and CRISPRCasFinder which do not support the search for such cases. We investigated the frequency of such cases in the comprehensive CRISPR array dataset collected with CRISPRidentify and found that 152 of the 39 378 identified arrays contained spacer deletion (see Supplementary Section 12).

### Tool availability

The CRISPRidentify is available as an open-source tool in GitHub https://github.com/BackofenLab/CRISPRidentify It is implemented and tested in Python version 3.7.6 and ML models exported following the model persistence from scikit-learn. https://scikit-learn.org/stable/modules/modelpersistence.html CRISPRidentify accepts a complete or partial genome sequences input and identifies all possible arrays using a different mode of setting.

## CONCLUSION

In this work, we present a pipeline for the detection of CRISPR arrays, CRISPRidentify, which includes a highly sensitive method for the detection of array candidates, followed by a data-driven, machine-learning based approach for the evaluation of these candidates. We demonstrate the ability of CRISPRidentify to accurately distinguish true CRISPR arrays from false ones. Although all existing CRISPR identification tools are highly sensitive, CRISPRidentify outperforms them by this measure, while also being substantially more specific. Using a DNA sequence as input, our pipeline performs three steps: (i) detection of repetitive elements and construction of array candidates, (ii) extraction of several array-related features and (iii) ML-based classification of array candidates based on the extracted features. The detected CRISPR arrays are then reported to the user accompanied by extensive annotation. In contrast to the existing tools, CRISPRidentify not only provides the user with the basic statistics of the identified CRISPR arrays, but also outputs a confidence score as an easily interpretable measure of the likelihood that a given genomic locus encompasses a CRISPR array. Thus, users can immediately assess the confidence of the identification of each potential CRISPR array, which is a substantial advantage over the other CRISPR detection tools. Another key feature of our approach is its flexibility. Whereas other tools use predetermined constant filtering criteria, CRISPRidentify dynamically builds such criteria based on the data employed during training. Thus, CRISPRidentify can adjust to new, previously unseen training data and use it to enhance the overall performance.

## SUPPLEMENTARY DATA

Supplementary Data are available at NAR Online.

## ACKNOWLEDGEMENTS

We thank the members of DFG SPP 2141 (‘Much more than Defence: the Multiple Functions and Facets of CRISPR–Cas’) for constructive discussions.

## REFERENCES

1. Barrangou, R. and van der Oost, J. (eds.) (2013) In: CRISPR–Cas Systems: RNA-mediated Adaptive Immunity in Bacteria and Archaea. Springer Press, Heidelburg, pp. 1–129.

2. Makarova, K.S., Wolf, Y.I., Alkhnbashi, O.S., Costa, F., Shah, S.A., Saunders, S.J., Barrangou, R., Brouns, S.J.J., Charpentier, E., Haft, D.H. et al. (2015) An updated evolutionary classification of CRISPR–Cas Systems. Nat. Rev. Microbiol., 13, 722–736.

3. Makarova, K.S., Wolf, Y.I., Iranzo, J., Shmakov, S., Alkhnbashi, O.S., Costa, F., Shah, S.A., Saunders, S.J., Barrangou, R., Brouns, S.J.J. et al. (2020) Evolutionary classification of CRISPRCas systems: a burst of class 2 and derived variants. Nat. Rev. Microbiol., 18, 67–83.

4. Levy, A., Goren, M.G., Yosef, I., Auster, O., Manor, M., Amitai, G., Edgar, R., Qimron, U. and Sorek, R. (2015) CRISPR adaptation biases explain preference for acquisition of foreign DNA. Nature, 520, 505–510.

5. Jackson, S.A., McKenzie, R.E., Fagerlund, R.D., Kieper, S.N., Fineran, P.C. and Brouns, S.J. (2017) CRISPR–Cas: adapting to change. Science, 356, eaal5056.

6. Amitai, G. and Sorek, R. (2016) CRISPR–Cas adaptation: insights into the mechanism of action. Nat. Rev. Microbiol., 14, 67–76.

7. Zhang, J., Rouillon, C., Kerou, M., Reeks, J., Brugger, K., Graham, S., Reimann, J., Cannone, G., Liu, H., Albers, S.-V. et al. (2012) Structure and mechanism of the CMR complex for CRISPR-mediated antiviral immunity. Mol. Cell, 45, 303–313.

8. Deng, L., Kenchappa, C.S., Peng, X., She, Q. and Garrett, R.A. (2012) Modulation of CRISPR locus transcription by the repeat-binding protein Cbp1 in Sulfolobus. NAR, 40, 2470–2480.

9. Shah, S.A., Hansen, N.R. and Garrett, R.A. (2009) Distribution of CRISPR spacer matches in viruses and plasmids of crenarchaeal acidothermophiles and implications for their inhibitory mechanism. Biochem. Soc. Trans., 37, 23–28.

10. Shah, S.A., Erdmann, S., Mojica, F.J. and Garrett, R.A. (2013) Protospacer recognition motifs: mixed identities and functional diversity. RNA Biol, 10, 891–899.

11. Leenay, R.T., Maksimchuk, K.R., Slotkowski, R.A., Agrawal, R.N., Gomaa, A.A., Briner, A.E., Barrangou, R. and Beisel, C.L. (2016) Identifying and visualizing functional PAM diversity across CRISPR–Cas systems. Mol. Cell, 62, 137–147.

12. Charpentier, E., Richter, H., van der Oost, J. and White, M.F. (2015) Biogenesis pathways of RNA guides in archaeal and bacterial CRISPR–Cas adaptive immunity. FEMS Microbiol. Rev., 39, 428–441.

13. Hille, F., Richter, H., Wong, S.P., Bratovič, M., Ressel, S. and Charpentier, E. (2018) The biology of CRISPR–Cas: Backward and forward. Cell, 172, 1239–1259.

14. Jackson, R.N. and Wiedenheft, B. (2015) A conserved structural chassis for mounting versatile CRISPR RNA-guided immune responses. Mol. Cell, 58, 722–728.

15. Haft, D.H., Selengut, J., Mongodin, E.F. and Nelson, K.E. (2005) A guild of 45 CRISPR-associated (Cas) protein families and multiple CRISPR/Cas subtypes exist in prokaryotic genomes. PLoS Comput. Biol., 1, e60.

16. Makarova, K.S., Grishin, N.V., Shabalina, S.A., Wolf, Y.I. and Koonin, E.V. (2006) A putative RNA-interference-based immune system in prokaryotes: computational analysis of the predicted enzymatic machinery, functional analogies with eukaryotic RNAi, and hypothetical mechanisms of action. Biol. Direct, 1, 7.

17. Makarova, K.S., Haft, D.H., Barrangou, R., Brouns, S.J.J., Charpentier, E., Horvath, P., Moineau, S., Mojica, F.J.M., Wolf, Y.I., Yakunin, A.F. et al. (2011) Evolution and classification of the CRISPR–Cas systems. Nat. Rev. Microbiol., 9, 467–477.

18. Shmakov, S., Aaron, A., Scott, D., Cox, D., Pyzocha, N., Yan, W., Abudayyeh, O.O., Gootenberg, J.S., Makarova, K.S., Wolf, Y.I. et al. (2017) Diversity and evolution of class 2 CRISPRCas systems. Nat. Rev. Microbiol., 15, 169–182.

19. Shah, S.A., Alkhnbashi, O.S., Behler, J., Han, W., She, Q., Hess, W.R., Garrett, R.A. and Backofen, R. (2019) Comprehensive search for accessory proteins encoded with archaeal and bacterial type III CRISPR-cas gene cassettes reveals 39 new cas gene families. RNA Biol., 16, 530–542.

20. Shmakov, S., Makarova, K.S., Wolf, Y.I., Severinov, K. and Koonin, E.V. (2018) Systematic prediction of genes functionally linked to CRISPR–Cas systems by gene neighborhood analysis. PNAS, 115, E5307–E5316.

21. Cass, S.D.B., Haas, K.A., Stoll, B., Alkhnbashi, O., Sharma, K., Urlaub, H., Backofen, R., Marchfelder, A. and Bolt, E.L. (2015) The role of Cas8 in type I CRISPR interference. Biosci. Rep., 35, e00197.

22. Brouns, S.J., Jore, M.M., Lundgren, M., Westra, E.R., Slijkhuis, R.J., Snijders, A.P., Dickman, M.J., Makarova, K.S., Koonin, E.V. and van der Oost, J. (2008) Small CRISPR RNAs guide antiviral defense in prokaryotes. Science, 321, 960–964.

23. Plagens, A., Richter, H., Charpentier, E. and Randau, L. (2015) DNA and RNA interference mechanisms by CRISPR–Cas surveillance complexes. FEMS Microbiol. Rev., 39, 442–463.

24. Kunin, V., Sorek, R. and Hugenholtz, P. (2007) Evolutionary conservation of sequence and secondary structures in CRISPR repeats. Genome Biol., 8, R61.

25. Horvath, P., Romero, D.A., Coûté-Monvoisin, A.C., Richards, M., Deveau, H., Moineau, S., Boyaval, P., Fremaux, C. and Barrangou, R. (2008) Diversity, activity, and evolution of CRISPR loci in Streptococcus thermophilus. J. Bacteriol., 190, 1401–1412.

26. Horvath, P. and Barrangou, R. (2010) CRISPR/Cas, the immune system of bacteria and archaea. Science, 327, 167–170.

27. Reimann, V., Alkhnbashi, O.S., Saunders, S.J., Scholz, I., Hein, S., Backofen, R. and Hess, W.R. (2016) Structural constraints and enzymatic promiscuity in the Cas6-dependent generation of crRNAs. Nucleic Acids Res., 45, 915–925.

28. Nickel, L., Ulbricht, A., Alkhnbashi, O.S., Frstner, K.U., Cassidy, L., Weidenbach, K., Backofen, R. and Schmitz, R.A. (2019) Cross-cleavage activity of Cas6b in crRNA processing of two different CRISPR–Cas systems in Methanosarcina mazei G1. RNA Biol., 16, 492–503.

29. Lange, S.J., Alkhnbashi, O.S., Rose, D., Will, S. and Backofen, R. (2013) CRISPRmap: an automated classification of repeat conservation in prokaryotic adaptive immune systems. Nucleic Acids Res., 41, 8034–8044.

30. Alkhnbashi, O.S., Costa, F., Shah, S.A., Garrett, R.A., Saunders, S.J. and Backofen, R. (2014) CRISPRstrand: predicting repeat orientations to determine the crRNA-Encoding strand at CRISPR loci. Bioinformatics, 30, i489–496.

31. Alkhnbashi, O.S., Meier, T., Mitrofanov, A., Backofen, R. and Voss, B. (2019) CRISPR–Cas bioinformatics. Methods, 172, 3–11.

32. Padilha, V.A., Alkhnbashi, O.S., Shah, S.A., De Carvalho, A.C. and Backofen, R. (2020) CRISPRCasIdentifier: Machine learning for accurate identification and classification of CRISPR-Cas systems. GigaScience, 9, giaa062.

33. Bland, C., Ramsey, T.L., Sabree, F., Lowe, M., Brown, K., Kyrpides, N.C. and Hugenholtz, P. (2007) CRISPR Recognition Tool (CRT): A tool for automatic detection of clustered regularly interspaced palindromic repeats. BMC Bioinformatics, 8, 209.

34. Couvin, D., Bernheim, A., Toffano-Nioche, C., Touchon, M., Michalik, J., Néron, B., Rocha, E.P.C., Vergnaud, G., Gautheret, D. and Pourcel, C. (2018) CRISPRCasFinder, an update of CRISRFinder, includes a portable version, enhanced performance and integrates search for Cas proteins. Nucleic Acids Res., 46, W246–W251.

35. Edgar, R.C. (2007) PILER-CR: fast and accurate identification of CRISPR repeats. BMC Bioinformatics, 8, 18.

36. Biswas, A., Staals, R.H.J., Morales, S.E., Fineran, P.C. and Brown, C.M. (2016) CRISPRDetect: a flexible algorithm to define CRISPR arrays. BMC Genomics, 17, i356–i356.

37. Abouelhoda, M., Kurtz, S. and Ohlebusch, E. (2004) Replacing suffix trees with enhanced suffix arrays. J. Discrete Algorithms, 2, 53–86.

38. Alkhnbashi, O.S., Shah, S.A., Garrett, R.A., Saunders, S.J., Costa, F. and Backofen, R. (2016) Characterizing leader sequences of CRISPR loci. Bioinformatics, 32, i576–i585.

39. Gudbergsdottir, S., Deng, L., Chen, Z., Jensen, J.V., Jensen, L.R., She, Q. and Garrett, R.A. (2011) Dynamic properties of the Sulfolobus CRISPR/Cas and CRISPR/Cmr systems when challenged with vector-borne viral and plasmid genes and protospacers. Mol. Microbiol., 79, 35–49.

40. He, J. and Deem, M.W. (2010) Heterogeneous diversity of spacers within CRISPR (clustered regularly interspaced short palindromic repeats). Phys. Rev. Lett., 105, 128102

41. Weinberger, A.D., Sun, C.L., Pluciński, M.M., Denef, V.J., Thomas, B.C., Horvath, P., Barrangou, R., Gilmore, M.S., Getz, W.M. and Banfield, J.F. (2012) Persisting viral sequences shape microbial CRISPR-based immunity. PLoS Comput. Biol., 8, e1002475.

42. Gesner, E.M., Schellenberg, M.J., Garside, E.L., George, M.M. and Macmillan, A.M. (2011) Recognition and maturation of effector RNAs in a CRISPR interference pathway. Nat. Struct. Mol. Biol., 18, 688–692.

43. Juranek, S., Eban, T., Altuvia, Y., Brown, M., Morozov, P., Tuschl, T. and Margalit, H. (2012) A genome-wide view of the expression and processing patterns of Thermus thermophilus HB8 CRISPR RNAs. RNA, 18, 783–794.

44. Sternberg, S.H., Haurwitz, R.E. and Doudna, J.A. (2012) Mechanism of substrate selection by a highly specific CRISPR endoribonuclease. RNA, 18, 661–672.

45. Lorenz, R., Bernhart, S.H., Honer Zu Siederdissen, C., Tafer, H., Flamm, C., Stadler, P.F. and Hofacker, I.L. (2011) ViennaRNA Package 2.0. Algorithms Mol Biol, 6, 26.

46. Altschul, S.F., Gish, W., Miller, W., Myers, E.W. and Lipman, D.J. (1990) Basic local alignment search tool. J. Mol. Biol., 215, 403–410.

47. Hyatt, D., Chen, G.L., Locascio, P.F., Land, M.L., Larimer, F.W. and Hauser, L.J. (2010) Prodigal: prokaryotic gene recognition and translation initiation site identification. BMC Bioinformatics, 11, 119.

48. Finn, R.D., Clements, J. and Eddy, S.R. (2011) HMMER web server: interactive sequence similarity searching. Nucleic Acids Res., 39, W29–W37.

49. Saeys, Y., Inza, I. and Larranaga, P. (2007) A review of feature selection techniques in bioinformatics. Bioinformatics, 23, 2507–2517.

50. Pedregosa, F., Varoquaux, G., Gramfort, A., Michel, V., Thirion, B., Grisel, O., Blondel, M., Prettenhofer, P., Weiss, R., Dubourg, V. et al. (2011) Scikit-learn: machine learning in Python. J. Mach. Learn. Res., 12, 2825–2830.

51. Zhang, Q. and Ye, Y. (2017). Not all predicted CRISPR–Cas systems are equal: isolated cas genes and classes of CRISPR like elements. BMC Bioinformatics, 18, 92.

52. Dosztányi, Z., Mészáros, B. and Simon, I. (2009) ANCHOR: web server for predicting.

53. Siguier, P., Gourbeyre, E. and Chandler, M. (2014) Bacterial insertion sequences: their genomic impact and diversity. FEMS Microbiol. Rev., 38, 865–891.

54. Enright, A.J., van Dongen, S. and Ouzounis, C.A. (2002) An efficient algorithm for large-scale detection of protein families. Nucleic Acids Res., 30, 1575–1584.

55. Urbanowicz, R.J., Olson, R.S., Schmitt, P., Meeker, M. and Moore, J.H. (2018) Benchmarking relief-based feature selection methods for bioinformatics data mining. J. Biomed. Inform., 85, 168–188.

56. Swarts, D.C., Mosterd, C., van Passel, M.W. and Brouns, S.J. (2012) CRISPR interference directs strand specific spacer acquisition. PLoS One, 7, e35888.

57. Yosef, I., Goren, M.G. and Qimron, U. (2012) Proteins and DNA elements essential for the CRISPR adaptation process in Escherichia coli. Nucleic Acids Res., 40, 5569–5576.

